# Structural models and considerations on the COA6, COX18 and COX20 factors that assist assembly of human cytochrome c oxidase subunit II

**DOI:** 10.1101/123349

**Authors:** Luciano A Abriata

**Affiliations:** Laboratory for Biomolecular Modeling, École Polytechnique Fédérale de Lausanne and Swiss Institute of Bioinformatics, 1015 Lausanne, Switzerland

**Keywords:** COXII, CuA assembly, copper homeostasis, molecular modeling, homology modeling, coevolution, I-TASSER, RaptorX

## Abstract

The soluble domain of cytochrome *c* oxidase subunit II (COX2), located in the outer side of the inner mitochondrial membrane, contains a binuclear copper site (CuA) through which electrons flow from cytochrome *c* to the core of the oxidase where oxygen reduction takes place. Being COX2 encoded in the mitochondrial genome, newly synthesized protein undergoes maturation steps in which it is translocated through and inserted into the inner mitochondrial membrane, and copper ions are loaded to form the CuA site. These steps are ensured by several protein factors in a complex pathway that is not fully understood, including copper-loading and disulfide-reduction proteins plus chaperones that assist proper membrane insertion. While the structure and function of copper-loading and disulfide-reducing proteins Sco1 and Sco2 have been quite studied at atomistic level, the latest biological studies have uncovered roles for other proteins that are not yet much understood at the structural level. In particular, recent experiments showed that membrane protein COX18 is a membrane-protein insertase for COX2, whereas membrane protein COX20 is a chaperone that stabilizes COX2 during translocation through the inner mitochondrial membrane, and soluble protein COA6 is part of the copper-loading pathway in conjunction with Sco1 and Sco2. This work reports structural models for COX18, COX20 and COA6, built judiciously from homology modeling, contact prediction-based modeling and transmembrane helix predictions, while considering the underlying biology. Implications and limitations of the models are discussed, and possible experimental routes to pursue are proposed. All models are provided as PyMOL sessions in the Supporting Information and can be visualized online at http://lucianoabriata.altervista.org/modelshome.html.

---

The binuclear copper site CuA located in the soluble domain of cytochrome *c* oxidase’s subunit II (COX2) is the entry port for electrons from cytochrome *c* to the oxidase, where oxygen reduction takes place at the end of the respiratory electron transport chain.^1^ Proper assembly of the multimeric cytochrome *c* oxidase in the inner mitochondrial membrane, including its several inorganic groups, is essential for cellular respiration, to an extent that defects result in serious energy-related diseases and syndromes.^2^ Therefore, assembly of the different proteinaceous and inorganic elements that make cytochrome *c* oxidase is ensured by a large array of proteins. The exact identities and functions of the proteins involved are still not clear, and we are far from a model explaining how the full oxidase is assembled. Recently, Bourens et al reported an important advancement in our understanding of how human COX2 is maturated, assigning specific roles to some of the protein factors involved in the process.^3^

COX2, as well as COX1 and COX3, the three core proteins that make every cytochrome *c* oxidase in all organisms, is encoded in the mitochondrial genome and is therefore synthesized in mitoribosomes. Human COX2 has two transmembrane (TM) helices, which get inserted in the inner mitochondrial membrane leaving the soluble CuA-containing domain in the intermembrane space (or periplasm in bacteria). The recent findings by Bourens et al combined with previous knowledge about COX2 maturation (refs. ^3–8^ and other works) lead to a model where cofactor COX20 stabilizes COX2 during insertion of its N-terminal TM helix, and cofactor COX18 promotes translocation of the full polypeptide (including the CuA ligands) across the inner mitochondrial membrane, being COX18 released upon subsequent binding of the SCO1-SCO2-COA6 copper-loading module to COX2-COX20 to finalize assembly of the CuA site.

Whereas there are structures available for human Sco1 and Sco2 in the Protein Data Bank, and human cytochrome *c* oxidase can be readily modeled through homology to the bovine protein, there are no clear sources of structural information for COA6, COX18 and COX20. This work reports structural models for these three proteins, and presents some structural considerations that could help drive future experiments and interpret existing data.

## Results and Discussion

*Transmembrane topology predictions to map orientations of COX18 and COX20 in the inner mitochondrial membrane*. It is first important to analyze possible TM topologies for the membrane-embedded proteins COX18 and COX20, by predicting TM helices and making sense of these predictions based on what is known about cytochrome *c* oxidase and its assembly process. A few different predictors return essentially the same predicted TM helices, so those of the the SPOCTOPUS server,^9^ one of the best performers in a recent assessment,^10^ are taken (Figure 1).

**Figure 1.**
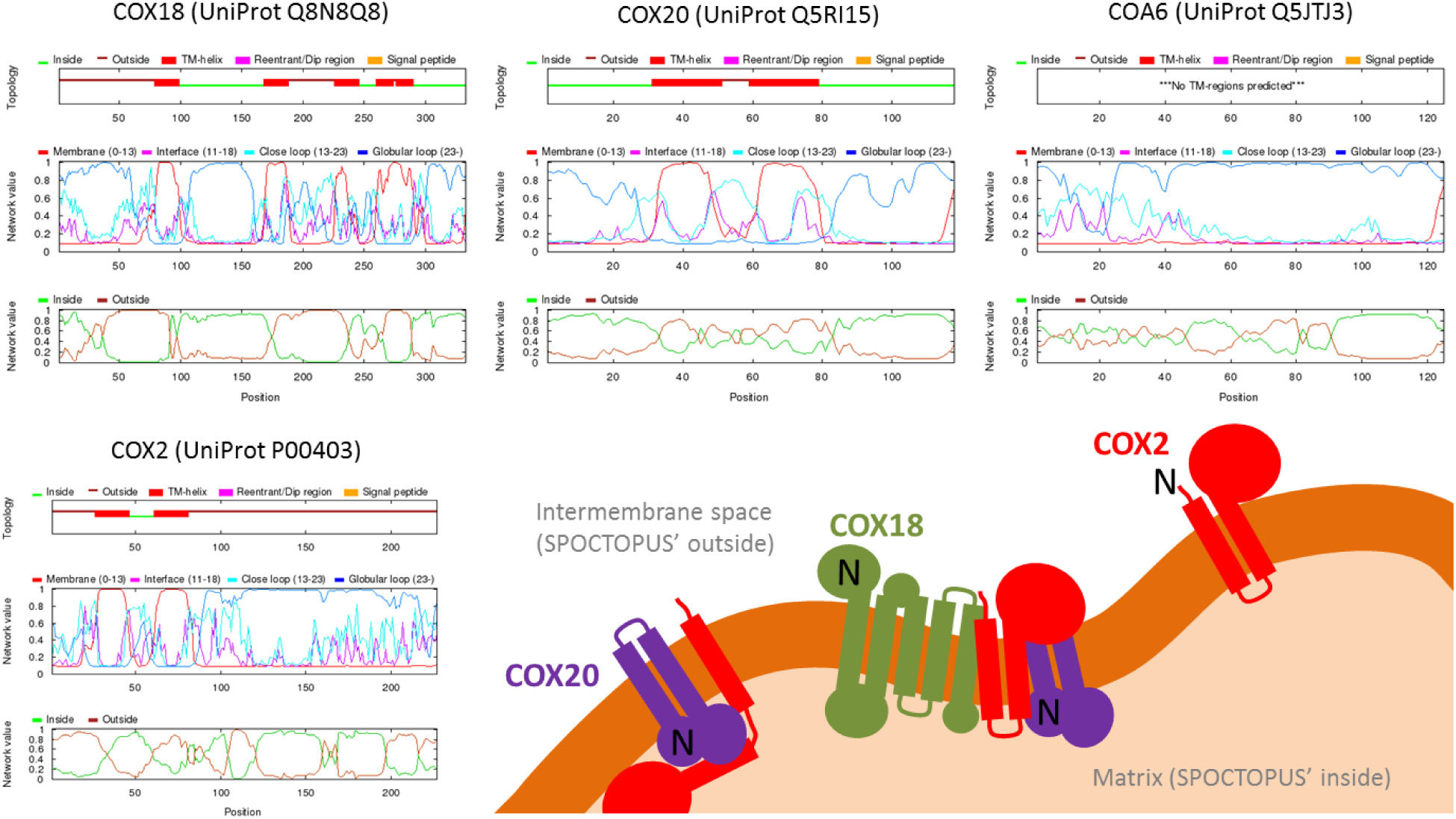
Prediction of TM helices by SPOCTOPUS for COX18, COX20, COA6 and COX2; and a model for COX2 maturation similar to that of Bourens et al accounting for predicted and known TM topologies.

SPOCTOPUS predicts COX18 and COX20 as TM proteins, as it predicts COX2, and COA6 as soluble. These predictions match the known localizations of the proteins and the scheme in Figure 7 of Bourens et al.^3^ However, matching the predicted insertion orientation for COX2 with its known orientation (N‐ and C-termini predicted outside, hence it corresponds to the intermembrane space), it turns out that COX20 is predicted to have an orientation opposite to that shown in that model. This implies COX20’s N‐ and C-termini being in the mitochondrial matrix, which makes sense as it would leave the large majority of its non-transmembrane domains in this space where the maturing COX2 needs to be stabilized.

### Strategies for modeling human COA6, COX18 and COX20

Sequences for human COA6, COX18 and COX20 were taken from UniProtKB entries Q5JTJ3, Q8N8Q8 and Q5RI15, respectively, choosing isoform 1 when more than one isoform was available. There are no clear templates for homology modeling of COA6, COX18 and COX20 from these sequences, other than moderate sequence similarity of COX18 to YidC membrane protein insertases according to SwissModel’s^11^ template detection module and as recognized by Bourens et al.^3^ Therefore two independent methods were used to build 3D models of these proteins, as in other similar studies:^12^ on one side, classical homology-based threading with the I-TASSER^13^ server, and on the other side modeling from contact predictions by the RaptorX server which analyzes large protein sequence alignments to estimate pairwise residue contacts and builds plausible models based on them.^14^ These two protocols are unrelated and use different data and methods; contact predictions rely on analysis of residue coevolution, a series of emerging methods for modeling without templates applicable even to disordered regions, alternative conformations, complexes and transmembrane domains.^14–22^

### Modeling of human COX18

For modeling COX18 through threading, I-TASSER selected PDB IDs 3WVF, 3WO6 and 3WO7 which correspond to crystal structures of YidC from *Escherichia coli* and *Bacillus halodurans*. The sequence-level match between COX18 and the possible templates is modest but enough for I-TASSER to estimate a template-modeling score^23,24^ of 0.63 for its model 1 (this scale ranges from 0 to 1; applied here higher scores suggest more reliable models).

The 5 models returned by I-TASSER are very similar to each other and to the templates between residues 40 and 295, out of 332 residues. The last 37 residues remain disordered with different conformations in the 5 models, most likely due to failure in threading through the template rather than reflecting true flexibility. In the 5 models, the first 40 residues adopt a conformation of a small alpha helix and two long β-sheets protruding out. This is an artifact that arises from an additional domain that *E. coli* YidC (PDB ID 3WVF) has relative to COX20, through which I-TASSER threaded this sequence segment.

RaptorX on COX18’s sequence returns 5 models that are very similar to each other in the sequence range from 80 to 295. The 5 RaptorX models are also very similar to the I-TASSER models and X-ray structures used for threading, except for some distortions and the orientation of the two polar helices that protrude out of the transmembrane core.

**Figure 2.**
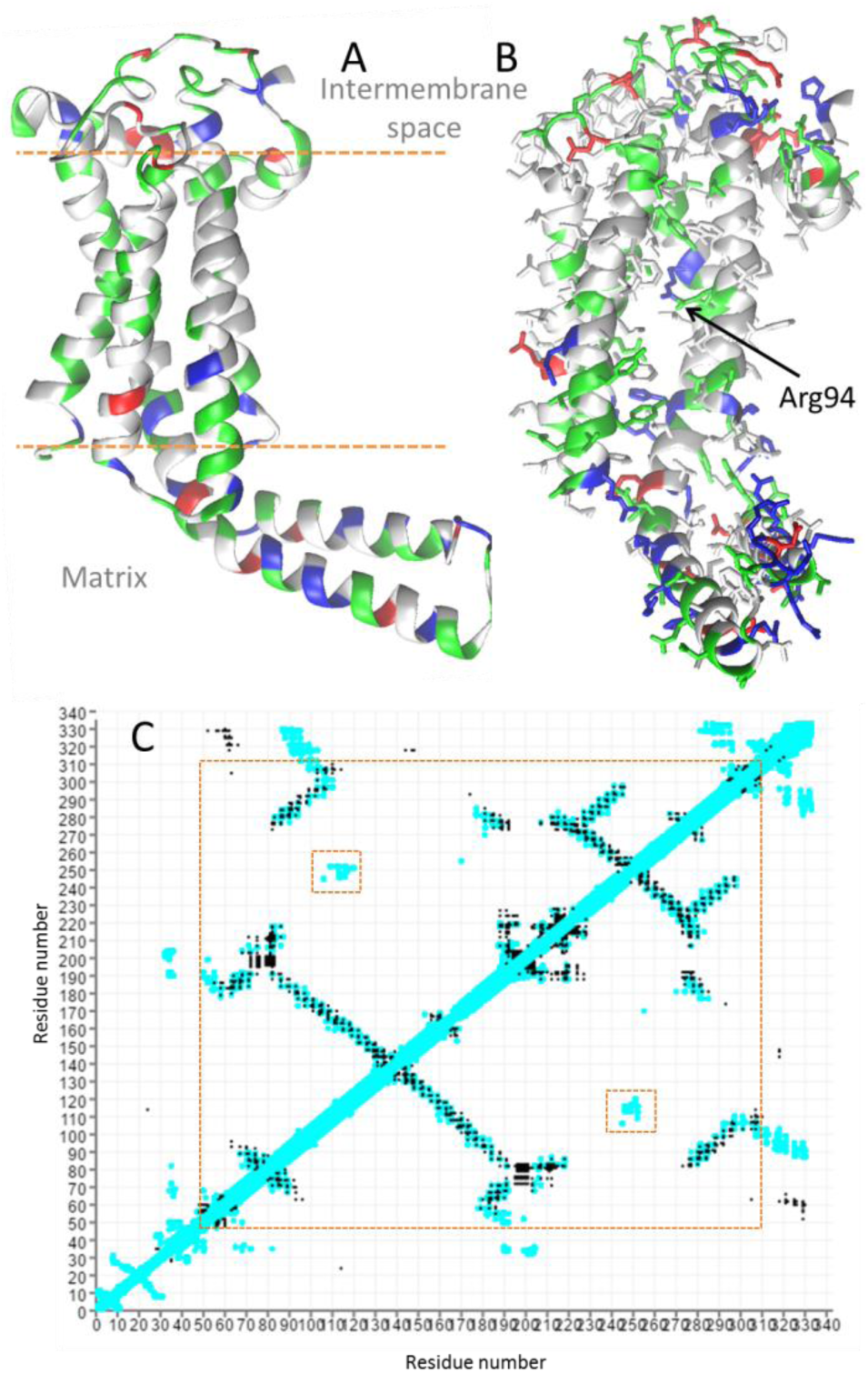
(A, B) I-TASSER model 1 for COX18 residues 50-300 with Arg94 indicated in (B). Amino acids colors: gray = hydrophobic, green = polar uncharged, blue = positive, red = negative. (C) Contact predictions from RaptorX (black dots) overlapped on the contact map of I-TASSER model 1 (cyan, 7Å cutoff distance between heavy atoms), performed with http://lucianoabriata.altervista.org/evocoupdisplay/raptorx.html. Internal red boxes point regions of no match.

Overall, based on these considerations and given the good match of RaptorX contact predictions to the I-TASSER models and the acceptable estimated TM-score of I-TASSER model 1, the latter provides a safe source of structural information for COX18 in the sequence segment from ∼50 to ∼300 (Figure 2). This model displays a shape and amino acid distribution consistent with the predicted TM topology, *i.e*. hydrophobic amino acid side chains exposed in the TM region but buried in the two helices that protrude into the mitochondrial matrix, plus aromatic and positively charged residues flanking the TM segments. The only positively charged residue that is buried inside the lumen formed by the TM helices is Arg94, which corresponds to a functional conserved site in membrane protein insertases that apparently “grabs” substrates to translocate them through the membrane.

### Modeling of human COA6

For modeling COA6 through threading, I-TASSER selected a series of PDB entries and chains that correspond to subunits 1 and 6B/12 of bovine heart cytochrome *c* oxidase. The sequence matches are moderate to weak, such that I-TASSER estimates a TM-score of 0.49 for its model 1.

The 5 models produced by I-TASSER are very similar to each other between residues ∼37 and ∼105, out of 125 total residues. Residues 105-125 and 1-50 are left disordered, most likely artificially *i.e*. due to bad template alignment. Within these regions modeled as disordered, residues 37-50 adopt very similar conformations.

Models built from contact predictions by RaptorX are all very similar to each other in the region from residue ∼50 to ∼113, and share a similar location of a loop spanning from residue ∼30 to ∼50 which adopts different disordered conformations across models. Overall, therefore, the I-TASSER and RaptorX models all suggest that residues ∼30-37 to 50 do adopt a structure, albeit probably not regular. However, the conformations of these loops are different in the I-TASSER and RaptorX models, resulting in different contact patterns (Figure 3A,B).

Since the conformations modeled by I-TASSER lack support from templates, in this case RaptorX models seem to provide safer sources of structural information for COA6 in the sequence segment from ∼30 to ∼113, with the caveat that segment ∼30-50 is poorly defined. It is important to note that running the DALI server^25^ on segment 30-113 of RaptorX’s model 1 returns some of the PDB entries that I-TASSER employed for modeling, highlighting that the similarity found at the sequence level holds at the structural level, even for models not built from homology-based protocols.

**Figure 3.**
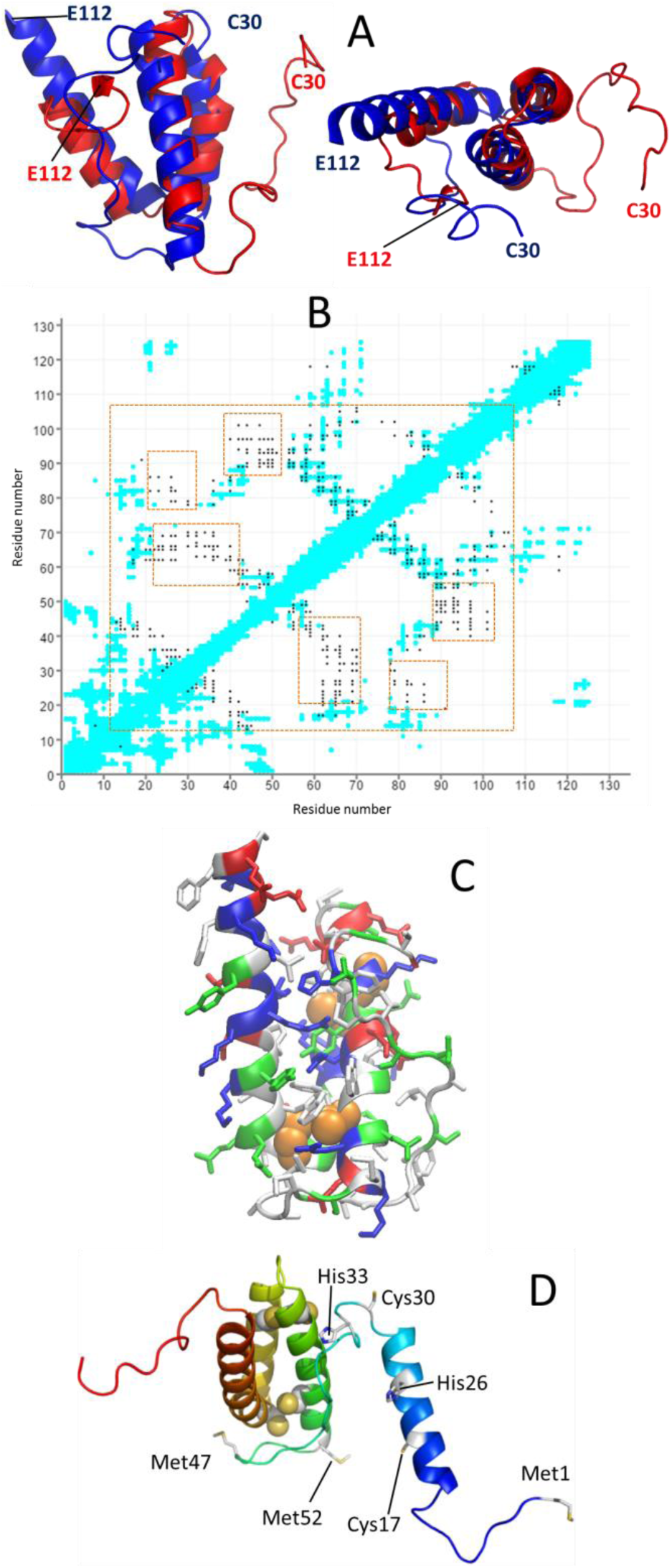
(A) Two views of I-TASSER (red) and RaptorX (blue) models #1 for COA6 30-112. (C) Contact predictions from RaptorX (black dots) overlapped on the contact map of I-TASSER model 1 (cyan, as in Figure 2), performed with http://lucianoabriata.altervista.org/evocoupdisplay/raptorx.html. Internal red boxes point regions of no match between contact maps. (C) RaptorX model 1 from 30 to 112, color-coded as in Figure 2B, with the four cysteines that likely form disulfide bonds as orange spheres. (D) Full RaptorX model 1, rainbow-colored from blue to red, with potential copper ligands shown as sticks and labeled.

In the I-TASSER models, two pairs of cysteines (58-90 and 68-79) are brought to close proximity, because analogous cysteine residues in the templates form disulfide bonds. The RaptorX models place the four conserved cysteins less closely than I-TASSER, but given their location and the low resolution of this kind of models, they might well form disulfide bonds. In such case, this leaves residues Cys17, His22, Cys30, His33, Met47 and Met52 as the potential ligands of the copper-binding site(s) required for COA6’s putative role in the maturation of the CuA site.

**Figure 4.**
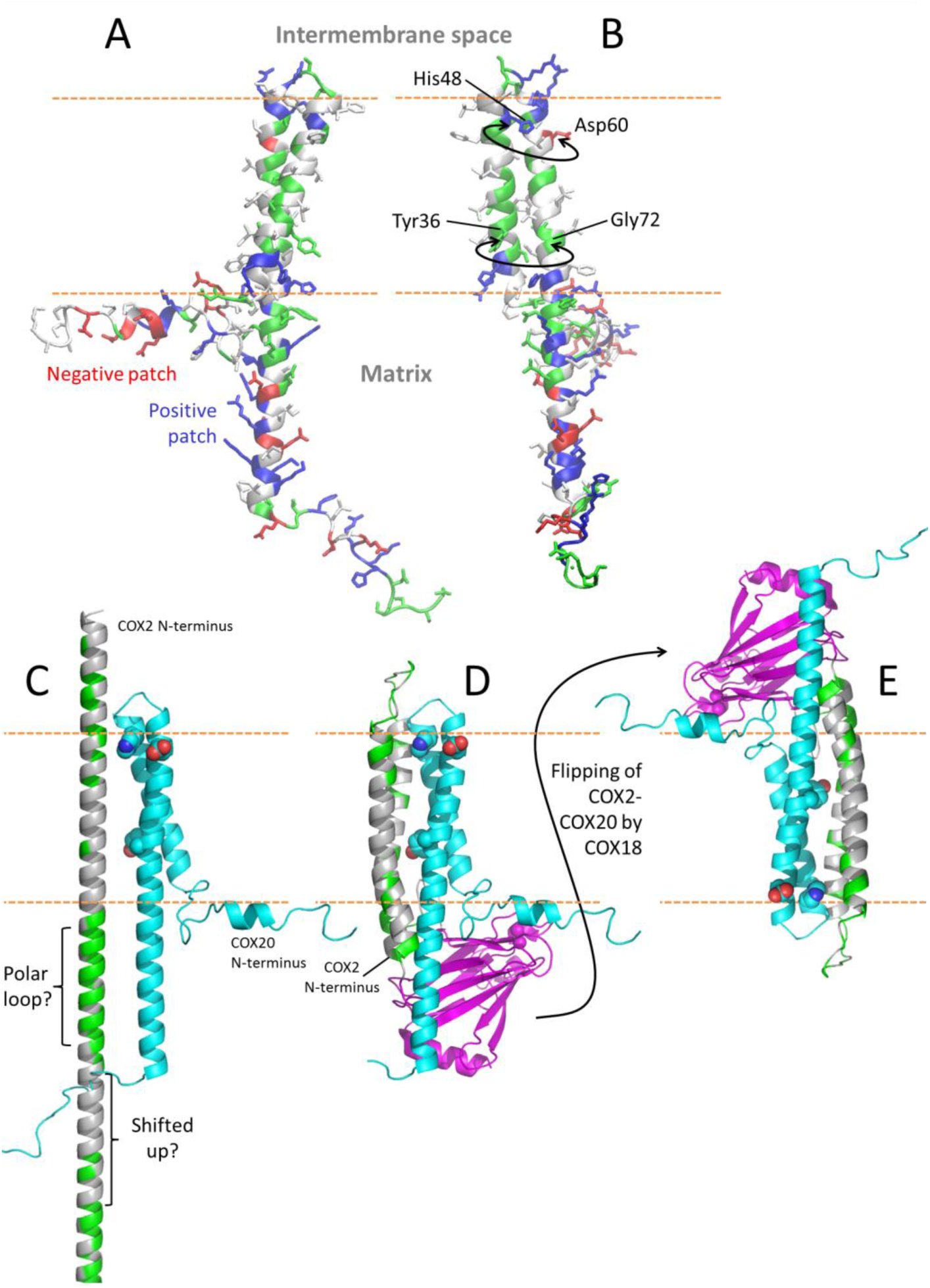
(A,B) Two views of the full RaptorX model 2 for COX20, showing in (B) the two pairs of residues that could establish contacts in a homodimer. Colors in this two panels are as in Figure 2A. (C-E) Possible arrangements that COX20 and COX2 could adopt if they interacted through the residues proposed to mediate COX20 homodimerization, when COX2 has been inserted into the inner mitochondrial membrane only through TM 1, assumed here to continue straight into TM 2 (C), or in a hypothetical situation where both TM1 and TM2 have been inserted but the whole COX2 has not yet been flipped to translocate its globular C-terminal domain (D) or has been flipped but is still not assembled to the rest of the growing oxidase (E). In panels C-E, the COX20 model (cyan) is RaptorX model 2, while COX2 (globular domain in magenta, TM region in gray and green for hydrophobic and hydrophilic residues, respectively. COX2 is chain B of PDB ID 3AG2 (bovine oxidase).

### Modeling of human COX20

For modeling COX20 through threading, I-TASSER selected disparate PDB entries, with each of the 10 alignment programs used by I-TASSER returning a different entry. This is because sequence matches are all weak, such that the estimated TM-score for model 1 is of only 0.34; therefore there is not much to profit from these models.

Models built from contact predictions by RaptorX display a long helix spanning residues ∼55-102, which interacts on its N-terminal half with a shorter helix spanning residues ∼30-50. The models are very similar between residues ∼30 and 102, out of 118 residues. The first 30 and last 16 residues are modeled as disordered and different to each other across the 5 RaptorX models. Part of this might reflect true flexibility, at least for segments 1-15 and 103-116 which are rich in small polar amino acids, prolines and charge repeats. Models 2 and 5 suggest a short alpha helix from 8 to 14, which given its amphipathic features and position could lie horizontally on the membrane.

DALI^25^ searches of these RaptorX models to the Protein Data Bank return several PDB entries, but their scores are bad and their annotations are not related to COX2 assembly pathways, copper handling nor chaperone functions that could imply structural and functional resemblance at null sequence identity.

Of the three proteins studied here, COX20 is clearly the hardest to model. However, the distribution of different types of amino acids in segment 30-102 of the RaptorX models is consistent with the independently predicted TM topology (Figure 4A). Namely, hydrophobic residues are placed with their side chains outside for helix segments 31-51 and 55-75, as expected for TM helices. Furthermore, aromatic and positively charged amino acids lie at the termini of the predicted TM segments. In turn, the lack of helix-disrupting amino acids suggests that the helical segment 56-102 would simply continue uninterruptedly from TM 55-75, possibly sticking out of the membrane into the mitochondrial matrix. Closer inspection suggests a possible dimerization interface in the TM region through pairs of residues Tyr36-Gly72 (the tyrosine OH often reaches glycine backbone oxygen to form a hydrogen bond) and His48-Asp60 (salt bridge or hydrogen bonds) (Figure 4B).

*Further discussion on the mechanisms by which COX18, COX20 and COA6 perform their functions, based on the structural models*. Membrane protein insertases related to COX18 have been quite studied through experiments, most notably YidC from *Bacillus halodurans*. Given the modeling results, it is likely that structure/function relationships known for YidC can be extrapolated to COX18, as commented above for the role of

Arg94.^26–28^ For COA6 and COX20, in contrast, the models introduced here are so far the only pieces of structural information available. Some more analyses beyond those presented together with the models follow.

COX20’s main role in COX2 maturation would be to stabilize it before translocation, when only its N-terminal TM helix (TM 1) is embedded in the inner mitochondrial membrane while its second TM helix (TM 2) and globular domain are still in the mitochondrial matrix. In this situation, one portion of COX2 that certainly requires stabilization is the very hydrophobic TM 2. Considering that the length of COX20’s C-terminal helix is roughly twice as long as that of a normal membrane-spanning helix, it is possible that this helix, protruding out of the membrane into the mitochondrial matrix, is meant to stabilize COX2’s TM helices when only TM 1 has been membrane-inserted. In such scenario (Figure 4C) COX2’s TM 1 and 2 would adopt a straight conformation, possibly with a loop in the connecting polar segment (“polar loop” in Figure 4C) that shifts TM 2 towards the membrane, so that both helical portions can interact with COX20’s long C-terminal helix through the membrane and into the mitochondrial matrix.

Another hypothetical situation is that COX20 can stabilize the two TM helices of fully membrane-inserted COX2 either before it is flipped to translocate its globular domain to the intermembrane space (Figure 4D) or after translocation but before being assembled into the growing oxidase (which would require transient flipping of COX20 when COX2 is flipped, Figure 4E).

COA6 has been implicated in copper loading to maturate the CuA site from the intermembrane space. It is in fact predicted soluble and the models present mostly polar residues on the surface. Portions of COA6’s N‐ and C-termini might be disordered, and the segment from residue ∼15 to ∼50 might form a loop for copper binding and transfer, as it contains amino acids often involved in copper binding:^29^ Cys17, His22, Cys30, His33, Met47 and Met52. Furthermore, according to the models, COA6 would have a small globular hydrophobic core. There is a hence reasonable chance that the protein be amenable to expression in soluble form for structural studies. Moreover, given its small size it might be amenable to NMR studies, which would allow for experimental exploration of its structure and dynamics, as well as its copper binding and transfer capabilities, as done on related systems.^30–32^

A peculiar observation about COA6 is that, as described in the modeling section above, I-TASSER detects weak sequence similarity to subunits of cytochrome *c* oxidase, and a DALI^25^ run on the models obtained from RaptorX contact predictions detects structural similarity to two subunits of cytochrome *c* oxidase: one of the matches is to a portion of subunit 1, and the other to COX subunit 6B/12. Interestingly, the latter docks on the soluble domain of COX2 in the full oxidase, an interaction that could mimic the native interaction that COA6 establishes with COX2 for copper delivery (Figure 5). Although in this pose COA6’s copper-binding loop does not seem to reach COX2’s CuA-binding loops, the hypothesis is worth exploring because COA6 and COX6B (and Sco2) have been shown to have overlapping roles in COX2 biogenesis.^33^

**Figure 5.**
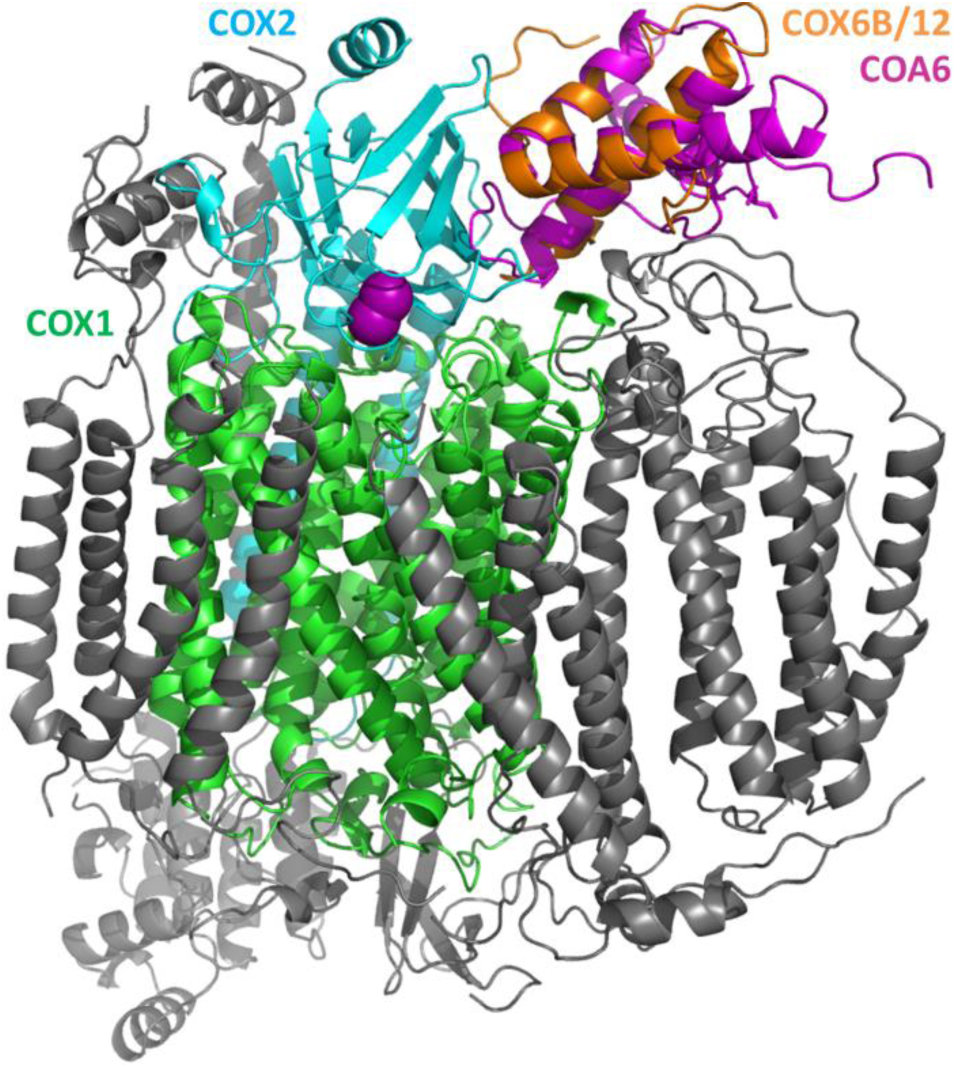
Bovine cytochrome *c* oxidase (PDB ID 1OCR) with subunits 1, 2 and 6B in green, cyan with magenta spheres on the CuA site, and orange, respectively, plus COA6 model 2 from RaptorX in magenta, aligned in 3D on COX6B/12.

## Conclusions

The models presented here hopefully contribute to understanding how COX2 assembly proceeds, either directly by comparing with available experimental data or by suggesting new experiments. In particular, mutations in COX20 could be designed to test dimerization and interactions with COX2; and *in vitro* studies on COA6 could be undertaken to get to know its structure, dynamics and copper binding and delivery functions.

## Methods

The RaptorX Contact Prediction and I-TASSER servers were run during March and April 2017. Overlays of contact predictions on model contact maps were drawn with the online tool at http://lucianoabriata.altervista.org/evocoupdisplay/raptorx.html. Figures were rendered with VMD^34^ and PyMOL.^35^

## Supporting Information

The Supporting Information material includes PyMOL sessions with RaptorX and I-TASSER models for COA6 and COX18, and RaptorX models for COX20; PyMOL sessions showing possible interactions of COX20 with COX2 as in Figure 4C and D, and a PyMOL session for the full oxidase structure 1OCR with COA6 RaptorX model 2 overlaid on COX 6B/12. Files are also available for download and online visualization at http://lucianoabriata.altervista.org/modelshome.html.

## Acknowledgements

I thank Dr. María Eugenia Zaballa (EPFL) for comments on the manuscript.

